# Analysis of immune subtypes across the epithelial-mesenchymal plasticity spectrum

**DOI:** 10.1101/2021.03.22.436535

**Authors:** Priyanka Chakraborty, Emily Chen, Isabelle McMullens, Andrew J. Armstrong, Mohit Kumar Jolly, Jason A. Somarelli

## Abstract

Epithelial-mesenchymal plasticity plays a critical role in many solid tumor types as a mediator of metastatic dissemination and treatment resistance. In addition, there is also a growing appreciation that the epithelial/mesenchymal status of a tumor plays a role in immune evasion and immune suppression. A deeper understanding of the immunological features of different tumor types has been facilitated by the availability of large gene expression datasets and the development of methods to deconvolute bulk RNA-Seq data. These resources have generated powerful new ways of characterizing tumors, including classification of immune subtypes based on differential expression of immunological genes. In the present work, we combine scoring algorithms to quantify epithelial-mesenchymal plasticity with immune subtype analysis to understand the relationship between epithelial plasticity and immune subtype across cancers. We find heterogeneity of epithelial-mesenchymal transition (EMT) status both within and between cancer types, with greater heterogeneity in the expression of EMT-related factors than of MET-related factors. We also find that specific immune subtypes have associated EMT scores and differential expression of immune checkpoint markers.

## Introduction

Epithelial-to-mesenchymal transition (EMT) is a cellular process in which epithelial cells lose epithelial characteristics, such as apical-basal polarity and tight cell-cell adhesions, and gain mesenchymal features, such as anterior-posterior polarization, focal cell contacts, and enhanced motility and invasiveness (Jolly et al. 2017; Nieto et al. 2016; J. Yang et al. 2020). Initially characterized in the field of embryology as a feature of normal development, EMT is now also known to play fundamental roles in multiple cellular processes, such as wound healing, fibrosis, and cancer (Stone et al. 2016). The regulation of EMT is complex and is controlled by the combined action of core EMT transcription factors (EMT-TFs, such as Snail/Slug, Twist1, and Zeb1/2) (Nieto et al. 2016; J. Yang et al. 2020), epithelial factors (such as GRHL2 (Somarelli et al. 2016; Gao et al. 2015) and OVOL1/2 (Roca et al. 2013)), post-transcriptional regulation, including microRNAs (Paterson et al. 2008; Gregory et al. 2008) and alternative splicing (Jolly et al. 2018; Bebee et al. 2015; Warzecha et al. 2009), epigenetic modifications (Skrypek et al. 2017), and post-translational regulation (Serrano-Gomez, Maziveyi, and Alahari 2016). In addition to these cell-intrinsic mechanisms, features of the tumor microenvironment, such as hypoxia (Saxena and Jolly 2019; Choi et al. 2017; Cannito et al. 2008; M.-H. Yang et al. 2008) and interactions between cancer cells and stromal (Y. Wang et al. 2021; L. Wang et al. 2018; M.-Q. Gao et al. 2010; Giannoni et al. 2010) and immune cells (Fedele and Melisi 2020; Romeo et al. 2019; Toh et al. 2011) also promote EMT.

Early work on EMT in cancer focused on EMT as a driver of metastatic dissemination. The reversion of cancer cells from a post-EMT-like state back to an epithelial-like phenotype by mesenchymal-to-epithelial transition (MET) was shown to be important for metastatic colonization subsequent to dissemination and seeding (Esposito et al. 2019;Somarelli et al. 2016; Stankic et al. 2013; Tsai et al. 2012; Ocaña et al. 2012). In addition to metastatic potential, however, EMT has also been shown to function in several other key cancer processes, including tumor-propagating/stemness-like phenotypes (Pasani, Sahoo, and Jolly 2020; Guen et al. 2017; Li et al. 2012; D. Guo et al. 2012; Wellner et al. 2009) and treatment resistance (Fischer et al. 2015; Zheng et al. 2015). Consistent with the role of EMT in metastatic dissemination and therapy resistance, the number and EMT phenotype of circulating tumor cells (CTCs) varies depending on a patient’s treatment response status, with treatment-refractory patients having more overall CTCs with a higher proportion of mesenchymal-type CTCs, and treatment responders having fewer overall CTCs with a higher proportion of epithelial-type CTCs (Yu et al. 2013).

Although early studies on EMT and MET viewed these phenotypic transitions as binary states, as this field of study has matured the simplified view of EMT/MET dynamics and roles in cancer progression has been shown to be more complex than originally described (Nieto et al. 2016; Biswas, Jolly, and Ghosh 2019). EMT, in some contexts, has now been shown to be dispensable for metastatic dissemination (Nieto et al. 2016; Zheng et al. 2015; Fischer et al. 2015). Similarly, MET has also been demonstrated to be dispensable for metastatic colonization, with two distinct paths to metastatic colonization (Somarelli et al. 2016; Jolly et al. 2017) and predicted by (Brabletz 2012). Likewise, while the importance of these processes is more complex, the dynamics of this gene regulatory program are also more nuanced than originally appreciated. Rather than a binary switch between states, EMT/MET is now viewed as a spectrum along which cells can have various hybrid E/M phenotypes (Nieto et al. 2016; Biswas, Jolly, and Ghosh 2019; Jolly et al. 2017). In fact, hybrid E/M phenotypes may be marks of increased cancer aggressiveness and metastatic ability (George et al. 2017). This has been observed not only in cancer cell lines, but also in clinical CTC samples (George et al. 2017; Armstrong et al. 2011; Yu et al. 2013).

Just as EMT promotes chemoresistance in multiple cancer types, the EMT/MET status of cancers has also been linked to resistance to novel immunotherapies. Immune checkpoint inhibitors have revolutionized the treatment landscape and vastly improved patient outcomes of several cancer subtypes, including non-small cell lung cancer (NSCLC) (Brahmer et al. 2018), melanoma (Weiss, Wolchok, and Sznol 2019), and renal cell carcinoma (RCC) (Weiss, Wolchok, and Sznol 2019; Rini et al. 2019). However, resistance to these agents is common, and there are several cancer subtypes that do not respond well to immune checkpoint blockade.

One possible explanation for poor immunotherapy response may be the relationship between the EMT status of a tumor and the expression of immune checkpoint molecules (Cao, Zhang, Kamimura, et al. 2011). For example, mesenchymal-like cancer cells have been shown to be capable of immunosuppression via interactions with stromal immune cells in the tumor microenvironment. In particular, Snail, a core EMT-TF, upregulates expression of CXCL2, a major mediator of myeloid-derived suppressor cell (MDSC) infiltration (Taki et al. 2018). MDSCs are major drivers of immunosuppression in the tumor microenvironment. MDSC infiltration renders cancer cells less susceptible to attack by CD8+ T cells and NK cells and leads to an increased ratio of immunosuppressive CD4+Foxp3+ Treg-like cells, which facilitates tumor growth (Qian et al. 2017). Snail1 and vimentin, a mesenchymal cytoskeletal marker, are positively correlated with programmed death-ligand 1 (PD-L1) scores (Kim et al. 2016). Furthermore, PD-L1 transcription is regulated by Snail1 and the miR200/ZEB1 axis (L. Chen et al. 2014). In contrast, E-cadherin (an epithelial marker) is negatively correlated with PD-L1 scores and epithelial-like cancer cells have higher numbers of infiltrating M1 macrophages and CD8+ T cells in the tumor microenvironment, which allows for greater susceptibility to immune checkpoint inhibitors (Dongre et al. 2017). It has also been proposed that the regulation between EMT and PD-L1 is actually bidirectional (Dongre et al. 2017; Jiang and Zhan 2020). This is supported by the observation that PD-L1 upregulates the EMT-TF, Twist (Cao, Zhang, Ritprajak, et al. 2011). In addition to this bidirectional regulation, there may be common inducers of EMT and immune evasion, including chronic inflammation, hypoxia, and metabolic reprogramming.

Our understanding of EMT, immune evasion, and other facets of cancer biology has been greatly aided by large, publicly-available gene expression datasets (Ding et al. 2018; Cancer Genome Atlas Research Network 2011). Analyses of these datasets have allowed researchers to characterize cancer types to an unprecedented level of detail. The coupling of these large genomics data sets with innovative algorithms that can uncouple bulk RNA-Seq data has also enabled inference of immune subtypes based on differential expression of immune-related genes within tumors (Eddy et al. 2020; B. Chen et al. 2018; Aran, Hu, and Butte 2017). These immune subtypes have prognostic value and can be used for survival stratification. Differences in outcome exist within and between cancers when stratified by immune subtypes.

In this study, we combined three novel EMT scoring metrics (Chakraborty et al. 2020) with immune subtype analysis from the iATLAS algorithm (Thorsson et al. 2018; Eddy et al. 2020) to understand the relationships between EMT and immune signature within and between cancer types. Our analyses demonstrate heterogeneity in both E/M phenotype and immune signature within a single cancer type. By comparing known drivers of epithelial and mesenchymal lineages across cancers, we reveal consistency in gene expression of epithelial factors across epithelial tumors, but cancer type-specific expression of a subset of mesenchymal drivers, suggesting that epithelial-derived cancers have convergence of key epithelial driver genes while mesenchymal-derived cancers may be driven by heterogeneous expression of one or more EMT-TFs. These drivers of E/M phenotype are also associated with distinct immune subtypes, with enrichment for specific immune subtypes across the EMT spectrum, illustrating the relationships between E/M status, immune subtype, with potential implications for patient response to immunotherapy.

## Results

### The EMT status of cancers is heterogeneous within and between cancer types

The EMT status of specific tumor types has been quantified using a variety of previously-established signatures, most of which use a limited set of molecules or functional traits and/or individual algorithms to measure the extent of EMT on a continuum (Mandal et al. 2016; Schliekelman et al. 2015; Ruscetti et al. 2016; Bhatia et al. 2019; S. Tripathi et al. 2020; Devaraj and Bose 2019; Puram et al. 2017). Here, to provide a robust comparison of EMT-like status across cancer types, we calculated the EMT scores with available RNA-Seq data from the Cancer Cell Encyclopedia (CCLE) using three distinct EMT scoring algorithms (George et al. 2017; Tan et al. 2014; Byers et al. 2013). Each of these three metrics – KS, MLR and GS76 - score the extent of EMT on a continuum, based on the expression of EMT-specific genes identified by various groups. These three methods use different gene lists and scoring methods: the GS76 method uses a weighted sum of expression levels of 76 genes, the KS method compares the cumulative distribution functions of epithelial and mesenchymal signatures, and the MLR method uses a multinomial logistic regression to calculate the probability of a sample to belong to varying EMT categories. KS and MLR score samples on a scale of [-1, 1] and [0, 2], respectively, while the GS76 metric has no pre-defined scale. Higher MLR or KS scores represent more mesenchymal samples while this is the inverse for GS76 scores (lower = more mesenchymal). Thus, KS and MLR scores of samples in a given dataset correlate positively with one another, and both KS and MLR correlate negatively with GS76 scores, as observed across multiple datasets (Chakraborty et al. 2020).

Consistent with their lineages of origin, carcinoma cell lines, such as colorectal, breast, stomach and prostate lines have lower median KS and MLR scores and higher GS76 scores as compared to mesenchymally-derived cancer cell lines, such as sarcoma, melanomas and glioma (**Figures 1A, S1A, S2A**). It is also noteworthy that the variance in EMT scores is higher in cell lines of epithelial lineages as compared to mesenchymally-derived cell lines (**Figures 1A, S1A, S2A**).

**Figure 1:**
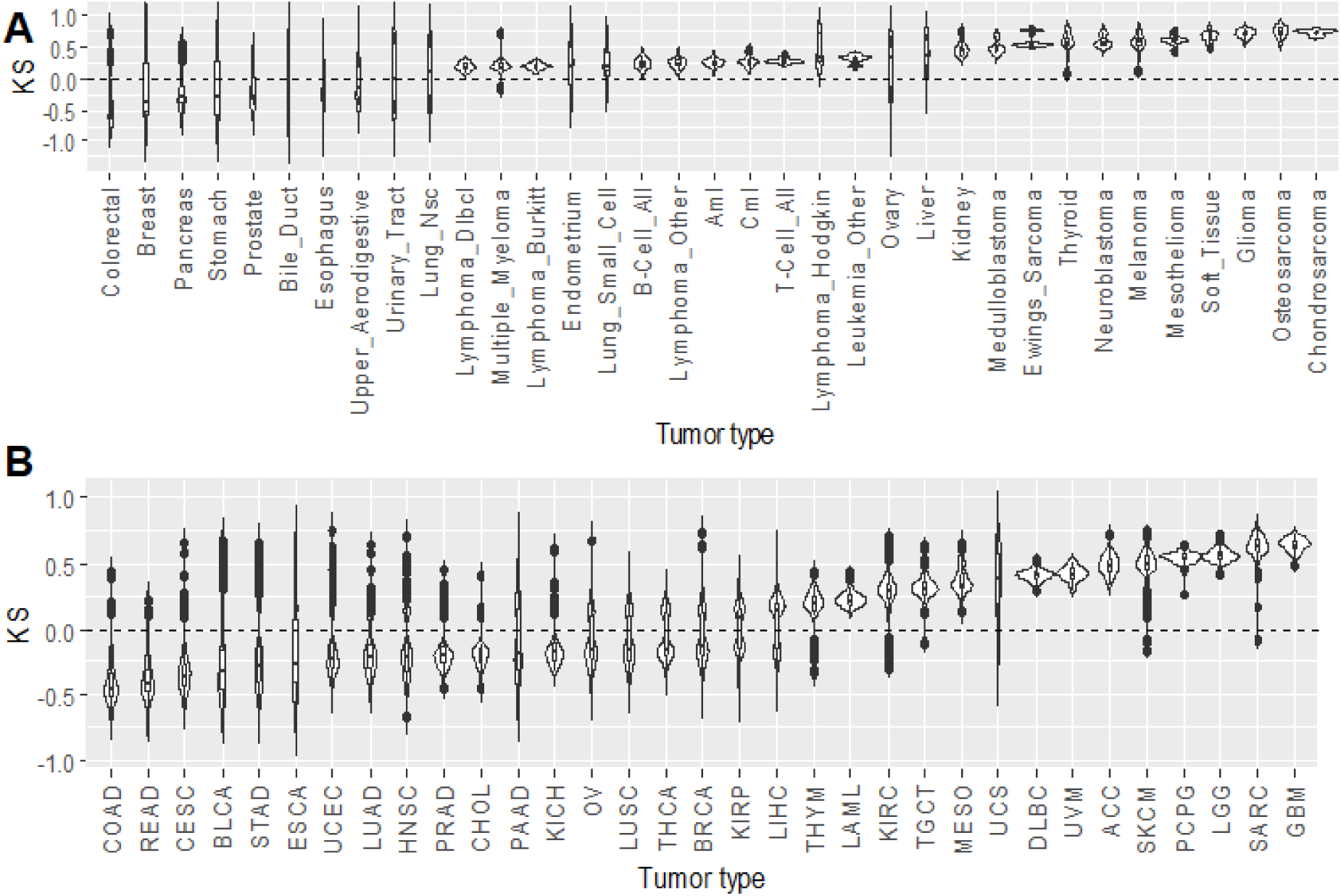
Carcinomas have more epithelial EMT scores. The KS algorithm scores samples on a scale of [-1, 1], with more positive scores representing more mesenchymal samples. **(A)** KS algorithm applied to RNA-Seq data from the Cancer Cell Encyclopedia (CCLE). **(B)** KS algorithm applied to RNA-Seq data from the Cancer Genome Atlas (TCGA); COAD = Colon adenocarcinoma, READ = Rectum adenocarcinoma, CESC = Cervical squamous cell carcinoma and endocervical adenocarcinoma, BLCA = Bladder urothelial carcinoma, STAD = stomach adenocarcinoma, ESCA = esophageal carcinoma, UCEC = uterine corpus endometrial carcinoma, LUAD = lung adenocarcinoma, HNSC = head and neck squamous cell carcinoma, PRAD = prostate adenocarcinoma, CHOL = cholangiocarcinoma, PAAD = pancreatic adenocarcinoma, KICH = kidney chromophobe, OV = ovarian serous cystadenocarcinoma, LUSC = lung squamous cell carcinoma, THCA = thyroid carcinoma, BRCA = breast invasive carcinoma, KIRP = kidney renal papillary cell carcinoma, LIHC = liver hepatocellular carcinoma, THYM = thymoma, LAML = acute myeloid leukemia, KIRC = kidney renal clear cell carcinoma, TGCT = testicular germ cell tumors, MESO = mesothelioma, UCS = uterine carcinosarcoma, DLBC = lymphoid neoplasm diffuse large B-cell lymphoma, UVM = uveal melanoma, SKCM = skin cutaneous melanoma, PCPG = pheochromocytoma and paraganglioma, LGG = brain lower grade glioma, SARC = sarcoma, GBM = glioblastoma multiforme.

We next investigated the distribution of EMT scores across tumor types from TCGA. Similar to the CCLE data, epithelial tumors (adenocarcinomas of the colon, rectum, stomach, prostate, etc.) have lower mean value of KS and MLR scores and a higher mean value GS76 score as compared to tumors derived from mesenchymal lineages (glioblastoma, glioma, sarcoma) (**Figures 1B, S1B, S2B**). Also consistent with CCLE data, mesenchymally-derived tumors have a lower variance in scores whereas carcinomas display higher variability in EMT score (**Figures 1B, S1B, S2B**), suggesting less heterogeneity in the EMT status of mesenchymally-derived cancers. To account for any potential bias in EMT score calculation that may be due to excess stromal contamination we applied a previously-reported combination of five tumor purity prediction methods (Aran, Sirota, and Butte 2015). EMT score was weakly correlated with tumor purity (**Figure S3A**). The ESTIMATE algorithm was the most highly correlated to EMT score across cancers, with correlations of −0.47, −0.23, and 0.17 to KS, MLR, and GS76, respectively (**Figure S3A**). Other metrics of tumor purity, such as immunohistochemistry and the LUMP, ABSOLUTE, and CPE algorithms, had a maximum correlation of −0.23 (CPE vs. KS; **Figure S3A**). We also re-analyzed TCGA samples in the 50th and 75th percentiles of tumor purity. While samples with higher tumor purity tended to have lower variance, samples with higher purity were not markedly different in EMT score compared to all samples in a given cancer type (**Figure S3B**). Together, these analyses suggest that, while single samples may be influenced to a moderate extent by tumor purity, EMT score for a given cancer type is driven predominantly by tumor cell lineage and not a consequence of high stromal contamination.

### EMT-TFs display heterogeneous patterns across cancers while MET factors are more highly correlated

Given the heterogeneity in EMT scores within and between carcinomas, we asked if this heterogeneity was correlated with differences in EMT-TFs and MET-associated factors. To address this question we assessed the levels of a panel of five well-studied EMT-inducing or EMT-associated factors – SNAI1, SNAI2, ZEB1, ZEB2, and TWIST1 (Subbalakshmi et al. 2021; J. Yang et al. 2004; Drápela et al. 2020; Vandewalle 2005) – and five MET-inducing or MET-associated factors – GRHL2, OVOL1, OVOL2, ESRP1, and ESRP2 (Saxena et al. 2020; Cieply et al. 2012; Jolly et al. 2018; Jeong et al. 2017; Fici et al. 2017) – across tumor types. From this analysis we noted the emergence of two large clusters: one set of tumors comprised predominantly of carcinomas, with higher levels of MET factors and low levels of EMT factors (top cluster, **Figure 2A**), and another set of tumors with the opposite trend, comprised mostly of mesenchymally-derived cancers (glioblastoma, glioma, and sarcoma). We also noted a distinct difference in the relationship between MET factors and EMT factors in these groups: In the carcinoma subset MET factors are all highly expressed, while this consistency across EMT factors is not observed in the predominantly-mesenchymal cancers. Instead, mesenchymal tumors are characterized by upregulation of one predominant EMT-factor, such as SNAI2 for uveal melanoma and TWIST1 for sarcoma (**Figure 2A**). Conversely, while the mesenchymal-like cluster display consistently low levels of MET factors, the epithelial-like cluster displays high expression of EMT factors in several cases. Principal component analysis indicated that expression of MET factors is the predominant contributor to the formation of the epithelial-like and mesenchymal-like clusters, while the EMT factors alone are unable to segregate clusters by epithelial/mesenchymal lineage (**Figure S4**).

**Figure 2:**
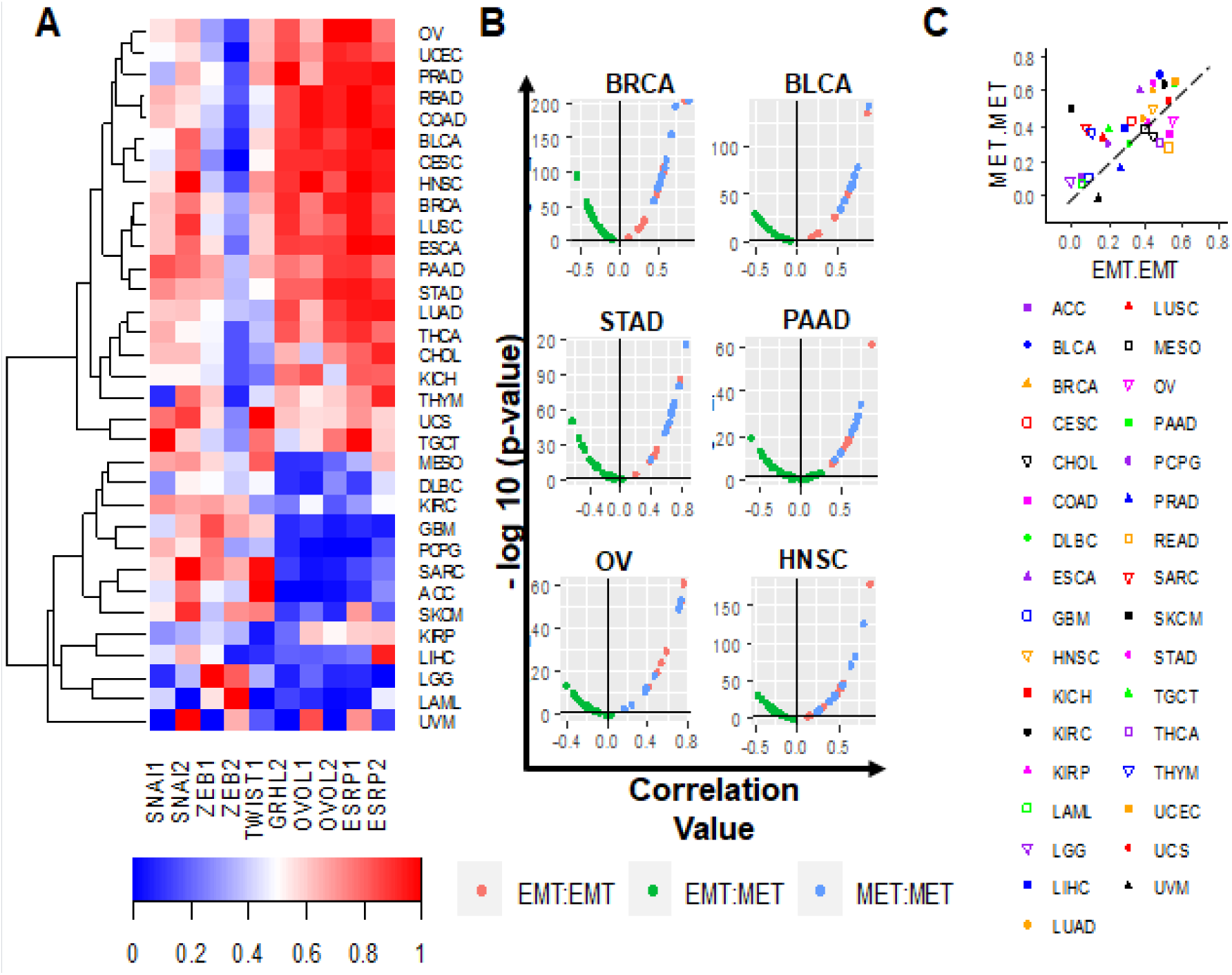
EMT factors are more heterogeneous across cancers than MET factors. EMT and MET marker expression across TCGA tumor types; COAD = Colon adenocarcinoma, READ = Rectum adenocarcinoma, CESC = Cervical squamous cell carcinoma and endocervical adenocarcinoma, BLCA = Bladder urothelial carcinoma, STAD = stomach adenocarcinoma, ESCA = esophageal carcinoma, UCEC = uterine corpus endometrial carcinoma, LUAD = lung adenocarcinoma, HNSC = head and neck squamous cell carcinoma, PRAD = prostate adenocarcinoma, CHOL = cholangiocarcinoma, PAAD = pancreatic adenocarcinoma, KICH = kidney chromophobe, OV = ovarian serous cystadenocarcinoma, LUSC = lung squamous cell carcinoma, THCA = thyroid carcinoma, BRCA = breast invasive carcinoma, KIRP = kidney renal papillary cell carcinoma, LIHC = liver hepatocellular carcinoma, THYM = thymoma, LAML = acute myeloid leukemia, KIRC = kidney renal clear cell carcinoma, TGCT = testicular germ cell tumors, MESO = mesothelioma, UCS = uterine carcinosarcoma, DLBC = lymphoid neoplasm diffuse large B-cell lymphoma, UVM = uveal melanoma, SKCM = skin cutaneous melanoma, PCPG = pheochromocytoma and paraganglioma, LGG = brain lower grade glioma, SARC = sarcoma, GBM = glioblastoma multiforme **(A)** Normalized EMT and MET marker expression across all TCGA tumor types. **(B)** Pairwise correlations between expression values of all EMT-EMT, MET-MET, and EMT-MET factor pairs across a subset of TCGA tumor types. **(C)** Plot of mean pairwise correlation coefficients for all EMT-EMT factor pairs (x-axis) versus mean pairwise correlation coefficients for all MET-MET factor pairs (y-axis) across all TCGA tumor types.

To further analyse this trend quantitatively, we calculated the pairwise correlations between expression values of all 10 EMT and MET factors across a subset of tumor types. Consistent with our qualitative observations above, the five MET factors are significantly positively correlated with each other, while the EMT and MET factors are significantly negatively correlated. However, unlike the MET factors, the correlation among different EMT factors is less consistent and often not statistically significant (**Figure 2B**). We also quantified the mean of the correlation coefficients for all pairwise correlations between any two EMT factors or for any two MET factors. These values are shown as x- and y-axes on a scatter plot, where each dot represents a tumor type. The majority of the tumors are above the x=y line, signifying that the average correlation between any two MET factors is greater than that between two EMT factors (**Figure 2C**). Similarly, visualization of the variance of the pairwise correlation coefficients for EMT and MET factors revealed a lower variability in MET factors as compared to EMT factors (**Figure S5**). These trends are also observed in CCLE cancer types (**Figures S6, S7**). Together, these analyses suggest that more heterogeneity in gene expression exists for EMT factors as compared to MET factors.

### EMT score is associated with specific immune subtypes in cancer

Activation of a partial or complete EMT has been associated with immune evasion across multiple cancers (S. C. Tripathi et al. 2016; Terry et al. 2017). A recent approach characterized immune tumor microenvironment across 33 cancer types in TCGA and defined six major immune subtypes spanning cancer types and molecular subtypes – C1 (wound healing), C2 (IFN-γ dominant), C3 (inflammatory), C4 (lymphocyte depleted), C5 (immunologically quiet), C6 (TGF-β dominant) (Thorsson et al. 2018). Given the relationship between EMT and immune checkpoint molecules (Terry et al. 2017; Kim et al. 2016; L. Chen et al. 2014) as well as immunomodulatory cytokines, we sought to understand the relationship between EMT status and immune subtype within and across cancers. To do this, we calculated EMT scores for all TCGA samples for which immune subtypes have been assigned. Overall, cancers with the C1 wound healing subtype and C2 IFN-γ dominant subtype tended to have more epithelial-like scores, while all other subtypes were more mesenchymal (**Figures 3A, S8A, C**). In particular, the C5 quiescence subtype is the most mesenchymal subtype (**Figure 3A-B, S8**). This observation is consistent with its sample composition, as C5 contained almost exclusively low-grade glioma samples (Thorsson et al. 2018). A leave-one-out analysis further demonstrated these trends, with the wound healing (C1) and IFN-γ (C2) signatures more epithelial than all other cancer samples, and the other immune signatures more mesenchymal than other signatures/samples (**Figures 3B, S8B, D**). We applied Singscore, a recently-developed rank-based single sample method to quantify the epithelial and mesenchymal scores separately (Foroutan et al. 2018) (**Figure 3C**). Singscore is a non-parametric method that calculates scores for ensembles of gene sets corresponding to epithelial and mesenchymal phenotypes. Using this method, the C5 samples showed high mesenchymal scores and very low epithelial scores. Interestingly, the mesenchymal scores for all six immune subtypes do not vary much while the epithelial score is the lowest for C5 (**Figure 3D**). The C6 TGF-β dominant immune subtype shows a high score for both the axes, which may suggest that cancers with this immune signature would have a more hybrid epithelial/mesenchymal (E/M) phenotype (**Figure 3D**).

**Figure 3:**
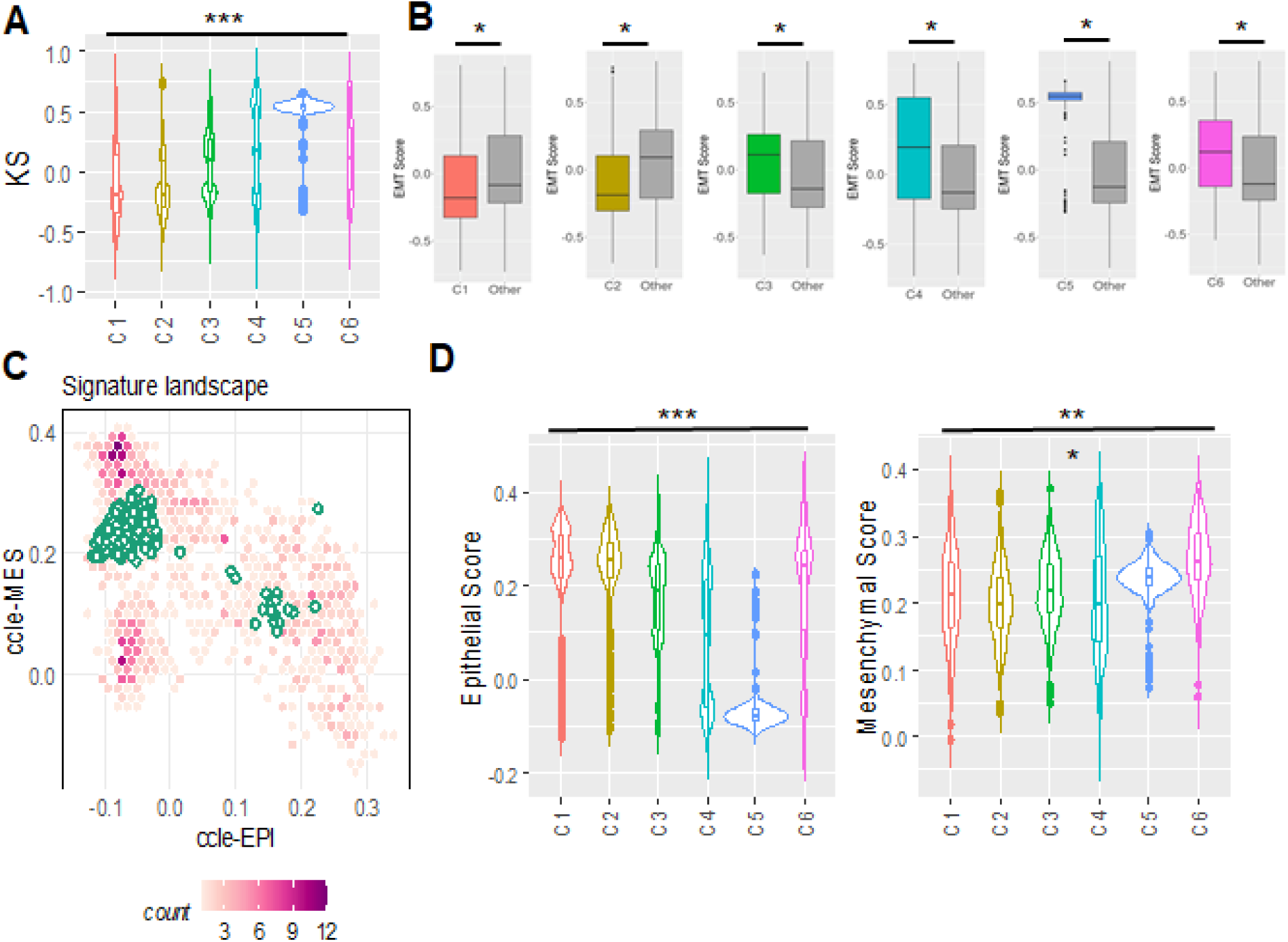
Immune subtypes are associated with EMT scores. EMT scores across TCGA immune subtypes; C1 = wound healing, C2 = IFN-γ dominant, C3 = inflammatory, C4 = lymphocyte depleted, C5 = immunologically quiet, C6 = TGF-β dominant **(A)** Plot of calculated KS score across all immune subtypes **(B)** Leave-one-out-analysis: pairwise comparison of each cancer immune subtype’s KS score to the KS scores of all other immune subtypes **(C)** Application of Singscore to C5 immune subtype samples to quantify epithelial and mesenchymal scores separately. **(D)** Singscore epithelial and mesenchymal scores across immune subtypes.

Given the marked heterogeneity in EMT scores and immune subtypes, we sought to understand whether particular cancer types are enriched in specific EMT scores and immune subtypes. Analysis of the upper and lower quartiles of EMT score across immune subtypes revealed distinct cancer types within each of the immune subtypes. For example, tumors in the upper quartile of the C1 wound healing subtype are enriched in colorectal cancer specimens as compared to the lower quartile of C1 (**Figure S9A**). Conversely, the composition of cancer types between upper quartile of EMT scores within the IFNγ-dominant C2 subtype are spread across multiple cancer types of diverse lineages, including, but not limited to, breast cancer, head and neck cancer, sarcomas, melanomas, and testicular germ cell tumors (**Figure S9B**). Likewise, the cancer types in upper and lower quartiles of EMT scores within the inflammatory (C3) subtype also differ substantially. More epithelial tumors within the C3 subtype (inflammatory) are enriched in renal clear cell carcinomas as compared to the lower quartile of C3 tumors, which is comprised of lung, prostate, breast, and thyroid cancers (**Figure S9A, B**).

To further understand how EMT and MET factors may be associated with specific immune signatures we performed separate analyses of EMT factors for each immune subtype. EMT factors Snail, Slug and Twist are most highly expressed in the C6 TGF-β dominant immune signature, with ZEB1 and ZEB2 most upregulated in the C5 immunologically quiescent signature (**Figure 4A-E**). Other immune subtypes have low to moderate levels of EMT factors, with wide distributions in expression across these subtypes. MET factors are predominantly expressed in C1-C3 signatures, with a bimodal distribution of expression in C4, low expression in C5, and relatively high expression in C6 (**Figure 4F-J**). Such co-expression of EMT and MET factors in C6 samples suggests that samples in the TGF-β dominant C6 immune subtype correspond to a more hybrid epithelial/ mesenchymal phenotype(s).

**Figure 4:**
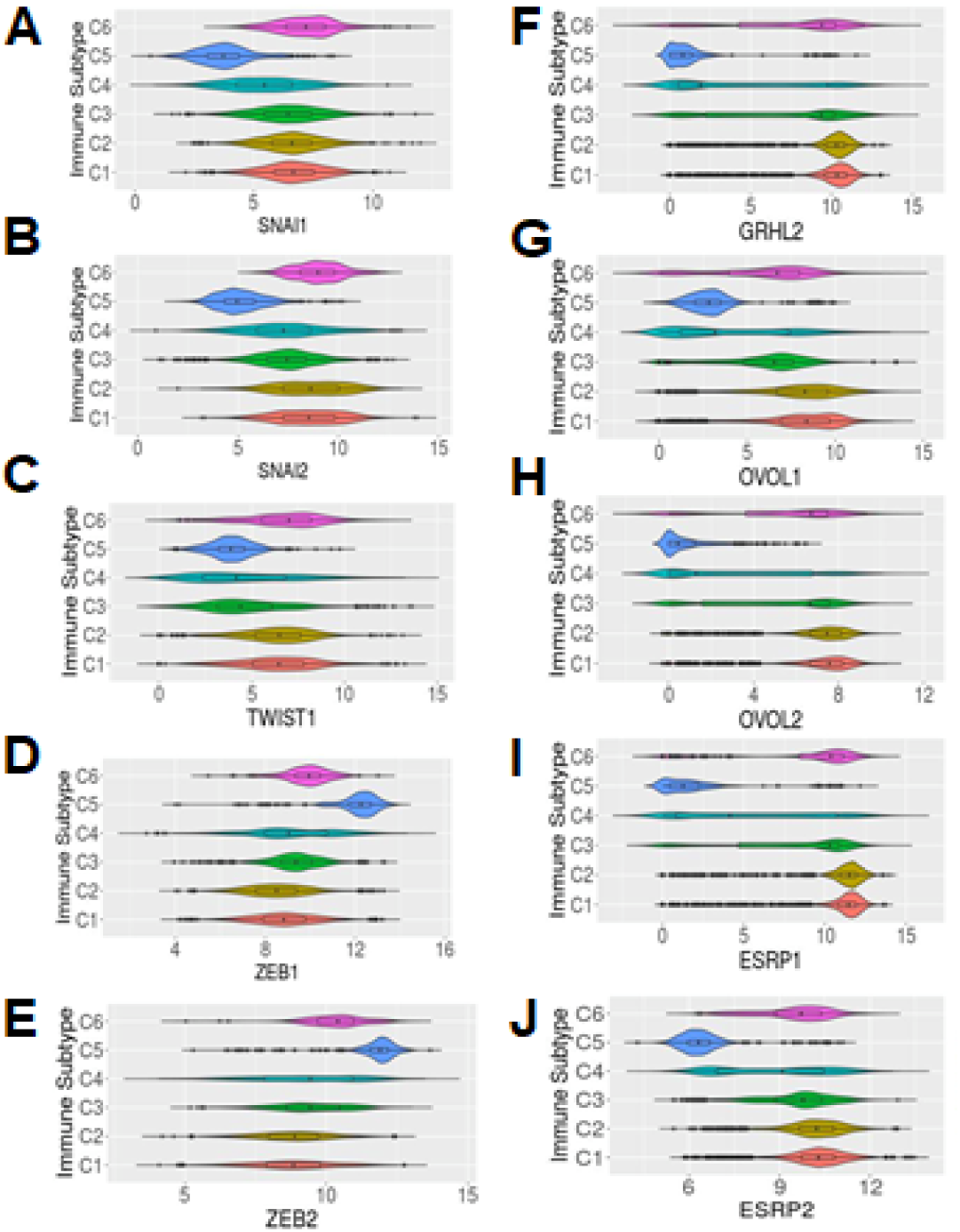
EMT and MET factor gene expression varies across immune subtypes. **(A-E)** Expression of EMT factors across immune subtypes; **(F-J)** Expression of MET factors acorss immune subtypes; **(A)** SNAI1, **(B)** SNAI2, **(C)** TWIST1, **(D)** ZEB1, **(E)** ZEB2, **(F)** GRHL2, **(G)** OVOL1, **(H)** OVOL2, **(I)** ESRP1, and **(J)** ESRP2 expression.

We also calculated the pairwise correlations between the EMT scores from the different scoring metrics (GS76, KS, MLR) and expression of EMT and MET factors from samples in each immune subtype. Across all immune signatures, these pairwise correlations showed consistent trends: GS76 scores correlated positively with levels of MET-factors and negatively with those of EMT-factors; KS and MLR scores followed the inverse trend (**Figure S10**). Among the EMT factors, ZEB1 and ZEB2 had the highest correlations with all EMT scoring metrics (**Fig S10**). The upregulation of ZEB1 and ZEB2 in C5, and the strong correlations of ZEB1 and ZEB with EMT scoring metrics further supports observations about the key roles of these two EMT-TFs in maintenance of a mesenchymal phenotype (Addison et al. 2021).

### Immune checkpoint markers across immune subtypes

We next investigated the levels of different immune checkpoint markers across the immune subtypes – CTLA-4 (cytotoxic T lymphocyte antigen-4), CD274 (PD-L1; programmed death-ligand 1) (Wei, Anang, and Sharma 2019), LAG3 (lymphocyte activation gene 3) (Solinas et al. 2019), CD276 (B7-H3) (S. Yang, Wei, and Zhao 2020), CD47 (cluster of differentiation 47) (Liu et al. 2015), and HAVCR2 (TIM-3) and its ligand LGASL9 (galectin 9) (Holderried et al. 2019) (**Figure 5A-G**). While expression of most immune checkpoint molecules varies across all immune subtypes, the immunologically quiescent C5 immune subtype has the lowest levels of CTLA-4, CD274, and CD276, but with HAVCR2 and CD47 expression similar to other immune subtypes (**Figure 5**). The TGF-β-dominant subtype (C6) displays elevated HAVCR2 and LGALS9 (**Figure 5**). The expression levels of these immune checkpoint markers are positively correlated with one another as well as with the single-sample GSEA scores for EMT and partial EMT (Puram et al. 2017) signatures (**Figure 5H**). These results suggest that common signalling pathways implicated in EMT may be associated with changes in levels of various immune checkpoint markers.

**Figure 5:**
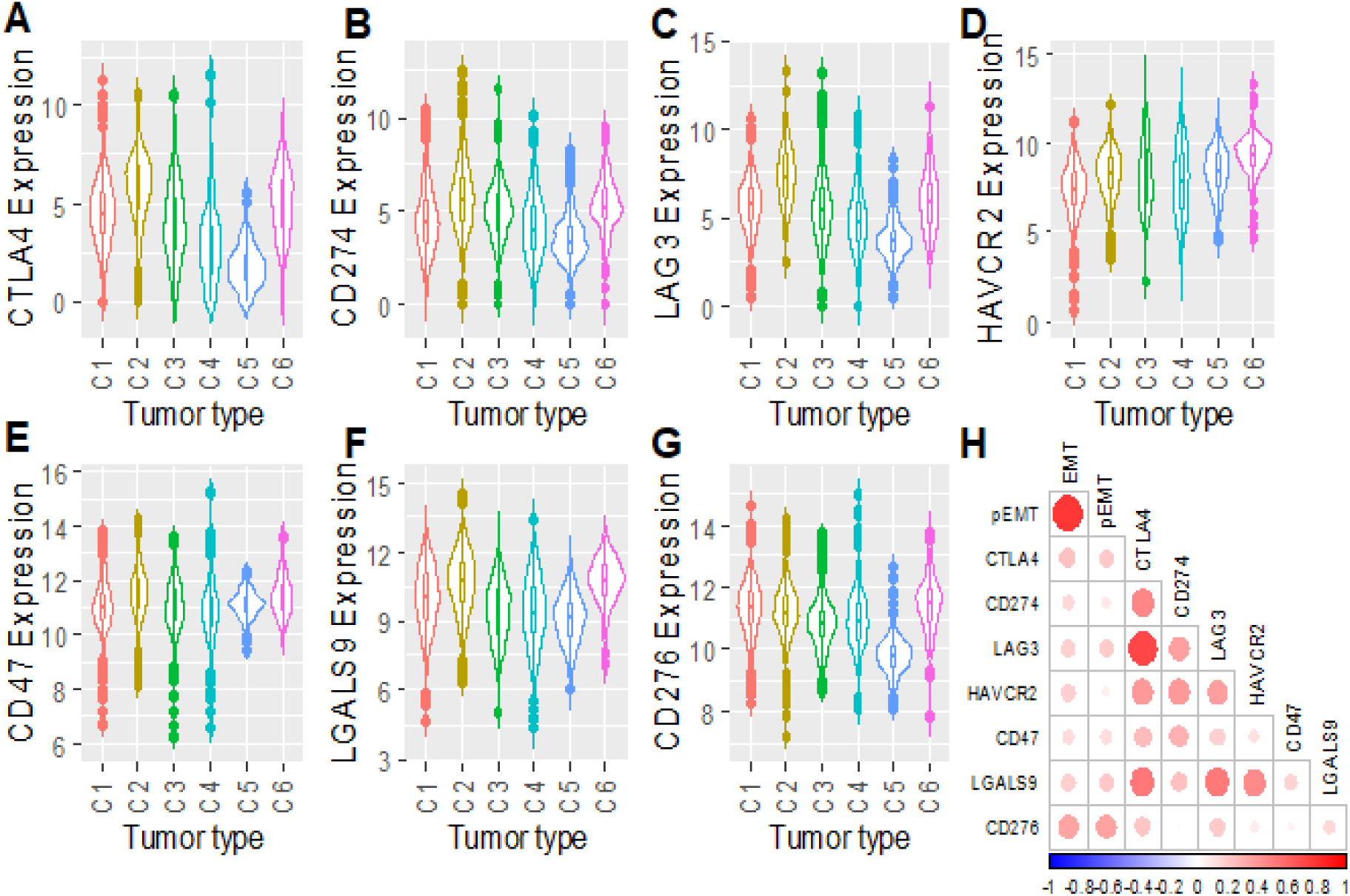
Immune checkpoint expression varies across immune subtypes. **(A)** CTLA4, **(B)** CD274, **(C)** LAG3, **(D)** HAVCR2, **(E)** CD47, **(F)** LGALS9, and **(G)** CD276 expression across immune subtypes. **(H)** Correlation matrix of EMT and pseudo-EMT scores with immune checkpoint markers.

## Discussion

Applying three distinct EMT scoring algorithms – KS, MLR, and GS76 – we characterized the diversity and heterogeneity of EMT status both within and across cancer types. These analyses revealed a higher variance in EMT score among cancers of an epithelial lineage as compared to mesenchymally-derived tumors. Similarly, analysis of EMT- and MET-associated factors revealed more heterogeneity in gene expression for EMT factors as compared to MET factors. While these data suggest that the EMT program is more heterogeneous than the MET program across cancers, the reasons for this are not clear. One possible explanation for this may be that the selected MET factors, such as GRHL2, OVOL1, and OVOL2 are indicative of epithelial lineages (Jolly et al. 2016; Hong et al. 2015; Lee et al. 2014), while the selected EMT factors may be more unique to a specific cellular lineage. It is possible that greater heterogeneity in the MET program would perhaps be more accurately reflected by analysing tissue-specific cadherins and keratins that mark specific lineages (Wahl and Spike 2017).

Another possible explanation may be that the MET program represents a more fixed, derived phenotype while the EMT program is more plastic in nature. This notion is supported by the observations that the EMT program is coupled to cancer stemness-like pathways (Mani et al. 2008; W. Guo et al. 2012; Wellner et al. 2009). Although it is possible that EMT scores may be skewed by differences in tumor:stromal ratios across samples and cancer types (Aran, Sirota, and Butte 2015; L. Wang et al. 2018), our analyses revealed low correlations between EMT score and tumor purity scores (Aran, Sirota, and Butte 2015), suggesting that the EMT scores were not substantially skewed by high stromal content. In addition, the scoring metrics were also consistent when applied to both TCGA and CCLE data, suggesting that these scoring metrics can be applied to both potentially-heterogeneous cancer samples and more homogeneous cancer cell lines. The remarkable heterogeneity in EMT and MET scores and EMT/MET factors across cancers underscores the concept of EMT/MET dynamics as a spectrum of phenotypic states (Nieto et al. 2016; J. Yang et al. 2020).

The present work sheds further light on our collective understanding of the potential cross-talk between cancer cells and immune subsets across the EMT spectrum. Both EMT scores and known EMT/MET factors were associated with certain immune subtypes. For example, cancers with C1 (wound healing) and C2 (IFN-γ dominant) subtypes tended to have more epithelial scores with high expression of MET factors, C5 (quiescent) was dominated by a high mesenchymal score with upregulation of ZEB1 and ZEB2, and C6 (TGF-β dominant) was enriched for a more hybrid epithelial/mesenchymal score. Consistent with this, the TGF-β dominant subtype shows upregulation of ZEB1 and ZEB2 as well as GRHL2, ESRP1, and ESRP2 (**Figure 4**). The hybrid E/M state plays an important role in therapeutic resistance and formation of highly tumorigenic CTC clusters (Jolly et al. 2016; Jolly 2015). TGF-β has been shown to activate ZEB1 through DNA methylation of the ZEB1 repressive microRNA, miR-200 (Gregory et al. 2011). Our prior work indicated that both GRHL2 and miR200s are necessary to induce MET in sarcomas(Somarelli et al. 2016), and it is possible that the TGF-β-driven subtype requires additional signals to drive the phenotype toward a more complete EMT. One limitation of this study is that the analysis of TCGA data was from almost exclusively primary tumor sites rather than metastases. Metastatic samples comprise just 3.4% of all TCGA solid tumor samples (https://portal.gdc.cancer.gov/). Given the considerable differences in EMT biology between primary tumors and metastases (Tsai et al. 2012; Ocaña et al. 2012; Bonnomet et al. 2012; Somarelli et al. 2016), it is likely that the biology of the immune subsets within metastatic microenvironments also differs.

Analysis of seven immune checkpoint molecules – CTLA-4, CD274/PD-L1, LAG3, CD276/B7-H3, CD47, HAVCR2/TIM-3, and LGAL29/galectin-9 – across the immune subtypes revealed the highest levels among C2 (IFN-γ dominant) and C6 (TGF-β dominant) subtypes and lowest in the C5 (quiescent) subtype. IFN-γ is a potent inducer of PD-L1 expression (Zou, Wolchok, and Chen 2016), which may explain the high levels of immune checkpoint molecules in the C2 subtype despite its more epithelial-like score. In addition, epithelial-like cancer cells are known to have higher numbers of infiltrating M1 macrophages and CD8+ T cells in the tumor microenvironment (Dongre et al. 2017), which is consistent with the observation that the C2 (IFN-γ dominant) subtype had the highest M1/M2 macrophage polarization and strong CD8 signal among all six subtypes (Thorsson et al. 2018). The TGF-β dominant subtype (C6) displayed increased levels of HAVCR2 (also known as Tim-3), and LGALS2 (galectin-9). Interestingly, HAVCR2 expression is upregulated by TGF-β (Yan et al. 2015; Wiener et al. 2007), and galectin-9 promotes TGF-β-induced signalling and subsequent conversion of CD4+CD25- T cells into regulatory T-cells(Lv et al. 2013; Wu et al. 2014). These observations suggest that patients enriched for the C6 subtype may benefit from TGF-β inhibitors in combination with HAVCR2 or galectin-9 suppression.

## Methods

All analyses were performed in R (version 3.4.4), and data was plotted using the ggplot2 package. Heatmaps were plotted using the gplots R package.

### TCGA datasets

TCGA gene expression datasets were obtained from https://xenabrowser.net/datapages/

### CCLE dataset

CCLE gene expression data was downloaded from https://portals.broadinstitute.org/ccle/data

### EMT scoring

Three different EMT scoring methods – KS, MLR, GS76 – were used to score samples separately in all the datasets as previously described (Chakraborty et al. 2020).

### ssGSEA analysis

ssGSEA analysis for various different gene sets were performed using GSVA R Bioconductor package with “ssgsea” option for method argument (Barbie et al. 2009).

### Statistical analysis

All the pairwise comparison significance was tested using student’s t-test and the multiple group comparisons significance was tested using ANOVA.

### Min**-**max standardization

The gene expression values and EMT scores were standardized in the range of 0 to 1 as following:

Where, *X_scaled_* is the min-max standardized value of a gene *X*.

### Principal component analysis

Principal component analysis was used to visualize the gene expression data of multiple variables (5 EMT and/or 5 MET factors). multidimensional gene expression and simulation data. To determine the correlation between variables and the representation of variables by the principal components, a correlation circle with squared cosines was plotted.

## Acknowledgments

JAS is supported by the Department of Defense (W81XWH-18-1-0189). AJA and JAS acknowledge funding support from NCI 1R01CA233585-02. The results published here are based, in part, upon data generated by the TCGA Research Network: https://www.cancer.gov/tcga.

## Supplementary figure legends

**Figure S1:**
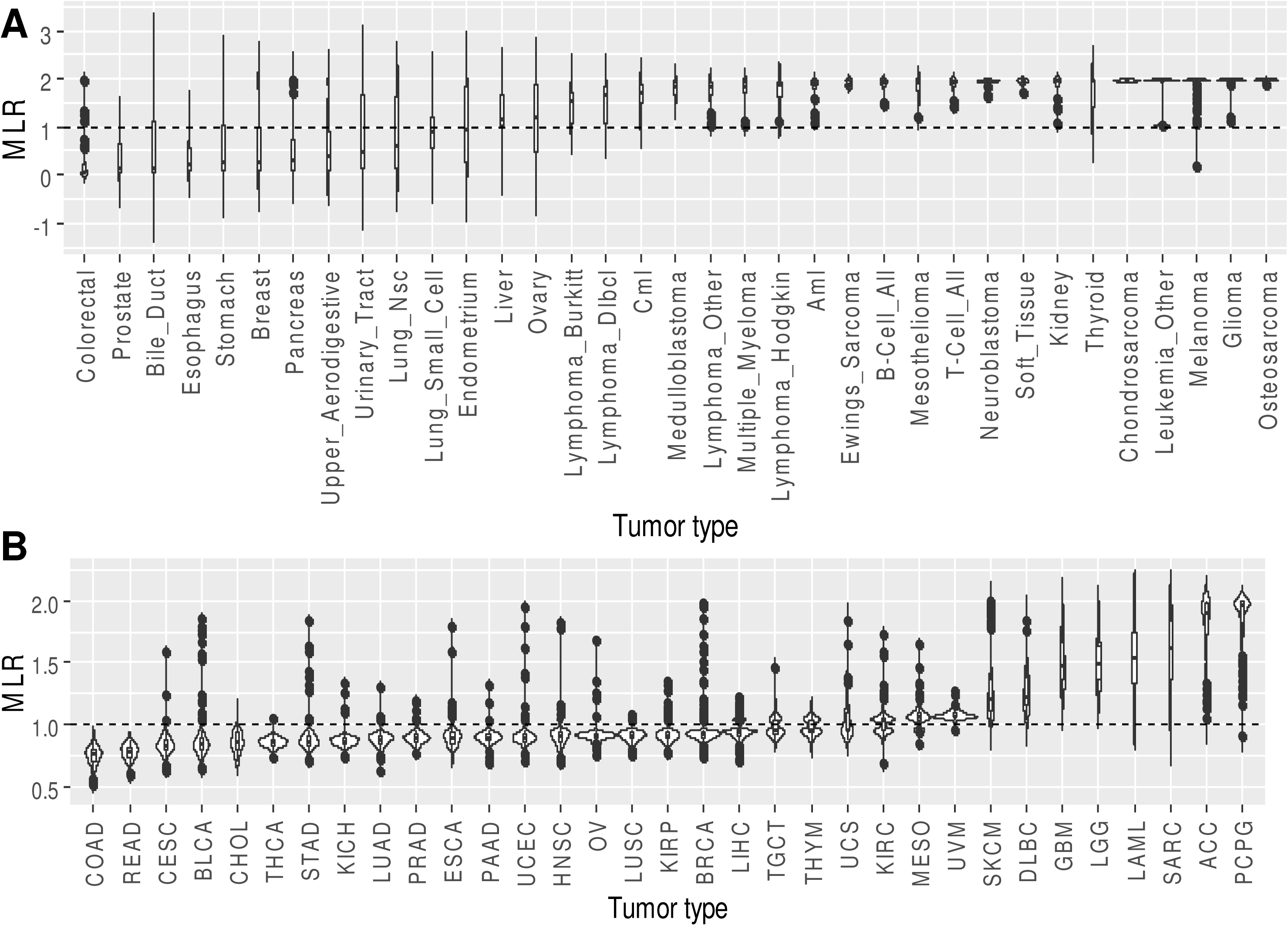
EMT scores across cancer types based on the MLR scoring metric. The MLR algorithm scores samples on a scale of [-1, 1], with more positive scores representing more mesenchymal samples. **(A)** MLR algorithm applied to RNA-Seq data from the Cancer Cell Encyclopedia (CCLE). **(B)** MLR algorithm applied to RNA-Seq data from the Cancer Genome Atlas (TCGA); COAD = Colon adenocarcinoma, READ = Rectum adenocarcinoma, CESC = Cervical squamous cell carcinoma and endocervical adenocarcinoma, BLCA = Bladder urothelial carcinoma, STAD = stomach adenocarcinoma, ESCA = esophageal carcinoma, UCEC = uterine corpus endometrial carcinoma, LUAD = lung adenocarcinoma, HNSC = head and neck squamous cell carcinoma, PRAD = prostate adenocarcinoma, CHOL = cholangiocarcinoma, PAAD = pancreatic adenocarcinoma, KICH = kidney chromophobe, OV = ovarian serous cystadenocarcinoma, LUSC = lung squamous cell carcinoma, THCA = thyroid carcinoma, BRCA = breast invasive carcinoma, KIRP = kidney renal papillary cell carcinoma, LIHC = liver hepatocellular carcinoma, THYM = thymoma, LAML = acute myeloid leukemia, KIRC = kidney renal clear cell carcinoma, TGCT = testicular germ cell tumors, MESO = mesothelioma, UCS = uterine carcinosarcoma, DLBC = lymphoid neoplasm diffuse large B-cell lymphoma, UVM = uveal melanoma, SKCM = skin cutaneous melanoma, PCPG = pheochromocytoma and paraganglioma, LGG = brain lower grade glioma, SARC = sarcoma, GBM = glioblastoma multiforme.

**Figure S2:**
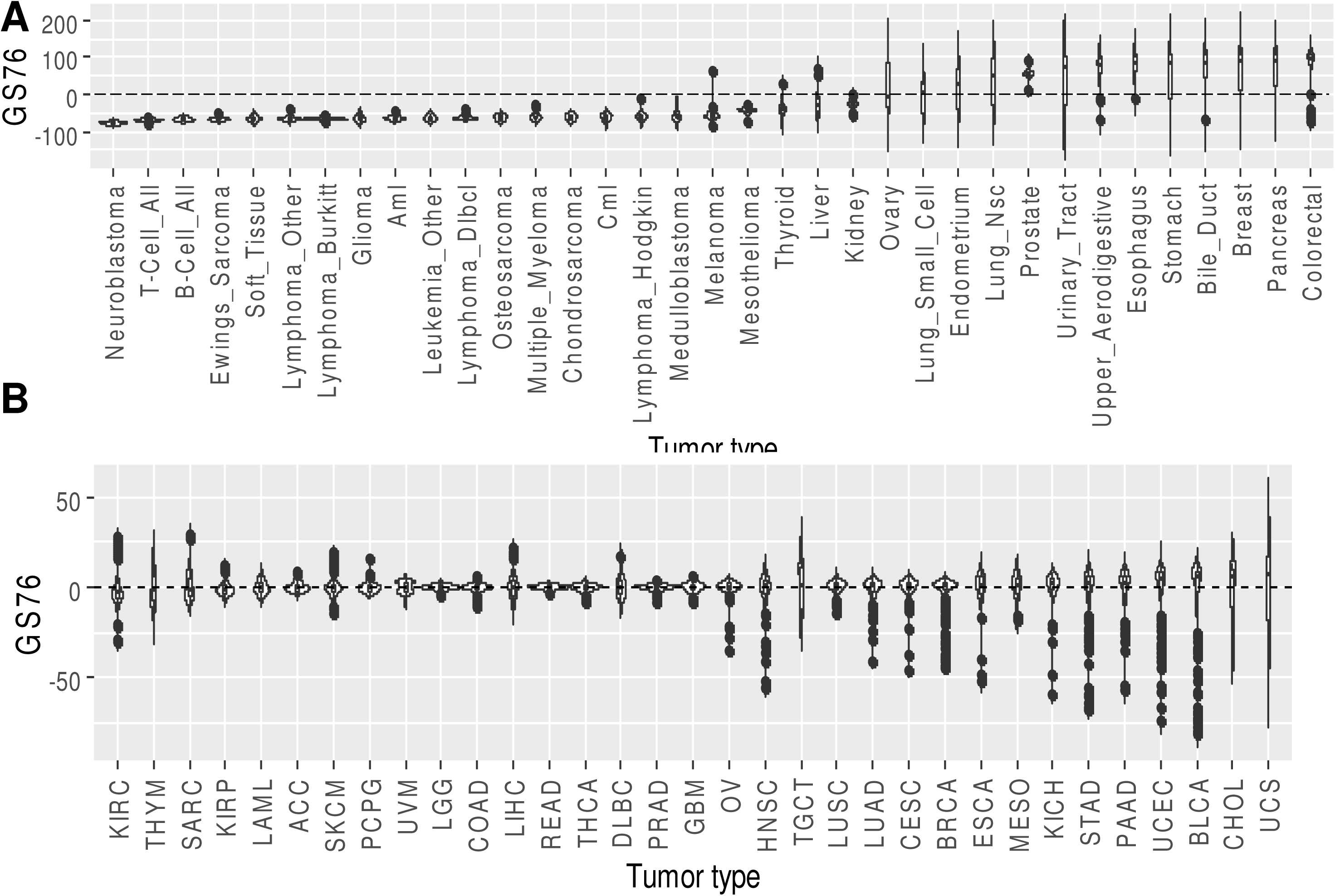
EMT scores across cancer types based on the 76GS scoring metric. There is no pre-defined scale for 76GS algorithm. More negative scores represent more mesenchymal samples. **(A)** 76GS algorithm applied to RNA-Seq data from the Cancer Cell Encyclopedia (CCLE). **(B)** 76GS algorithm applied to RNA-Seq data from the Cancer Genome Atlas (TCGA); COAD = Colon adenocarcinoma, READ = Rectum adenocarcinoma, CESC = Cervical squamous cell carcinoma and endocervical adenocarcinoma, BLCA = Bladder urothelial carcinoma, STAD = stomach adenocarcinoma, ESCA = esophageal carcinoma, UCEC = uterine corpus endometrial carcinoma, LUAD = lung adenocarcinoma, HNSC = head and neck squamous cell carcinoma, PRAD = prostate adenocarcinoma, CHOL = cholangiocarcinoma, PAAD = pancreatic adenocarcinoma, KICH = kidney chromophobe, OV = ovarian serous cystadenocarcinoma, LUSC = lung squamous cell carcinoma, THCA = thyroid carcinoma, BRCA = breast invasive carcinoma, KIRP = kidney renal papillary cell carcinoma, LIHC = liver hepatocellular carcinoma, THYM = thymoma, LAML = acute myeloid leukemia, KIRC = kidney renal clear cell carcinoma, TGCT = testicular germ cell tumors, MESO = mesothelioma, UCS = uterine carcinosarcoma, DLBC = lymphoid neoplasm diffuse large B-cell lymphoma, UVM = uveal melanoma, SKCM = skin cutaneous melanoma, PCPG = pheochromocytoma and paraganglioma, LGG = brain lower grade glioma, SARC = sarcoma, GBM = glioblastoma multiforme.

**Figure S3:**
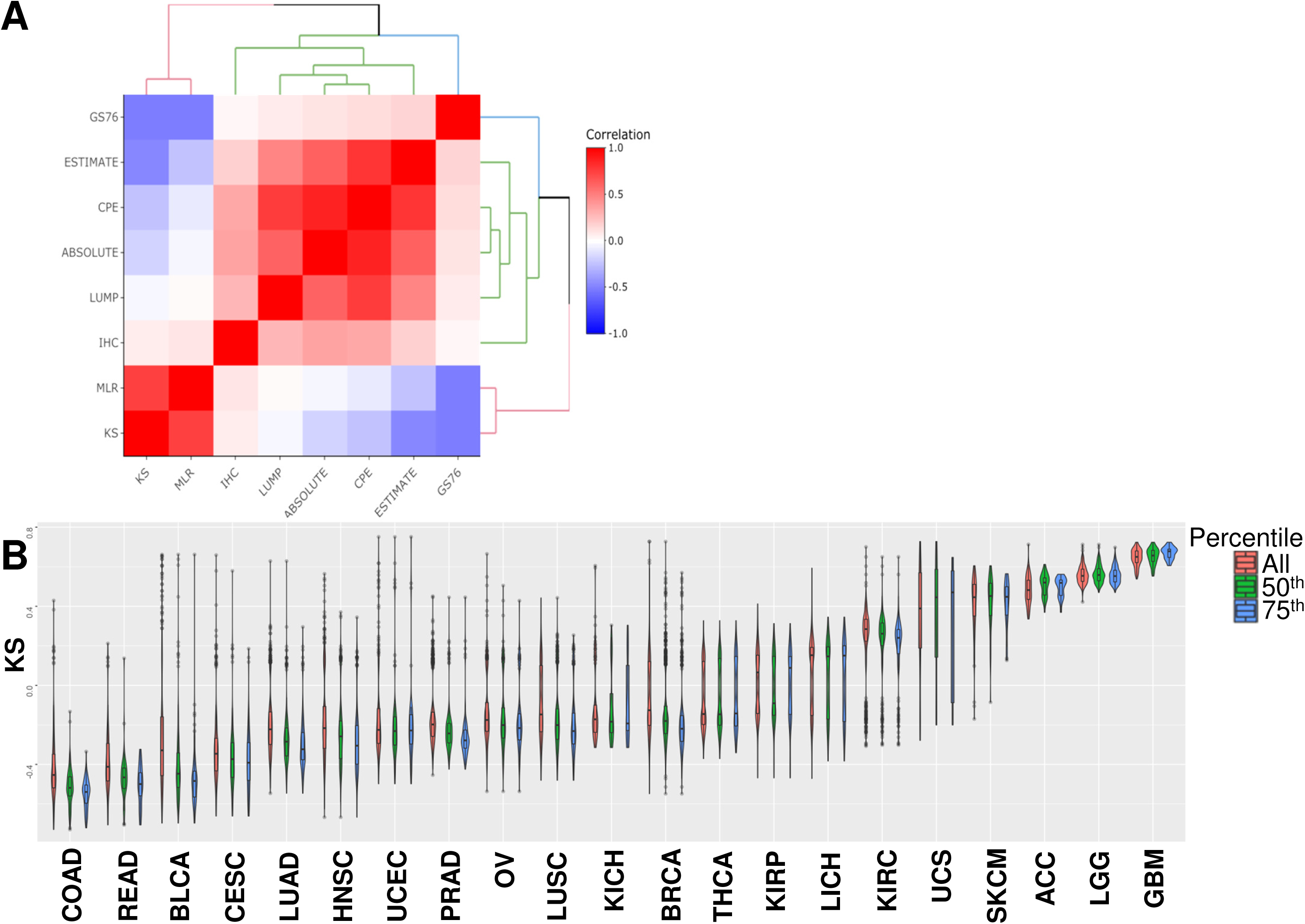
Correlation between EMT score and tumor purity. **(A)** Four metrics of tumor purity, including immunohistochemistry (IHC) and the LUMP, ABSOLUTE, and CPE algorithms, correlated to differing degrees with EMT scores calculated by the KS, MLR, and GS76 algorithms. (B) KS-based EMT scores with all samples, top 50^th^ percentile of tumor purity score, and top 75^th^ percentile of tumor purity score.

**Figure S4:**
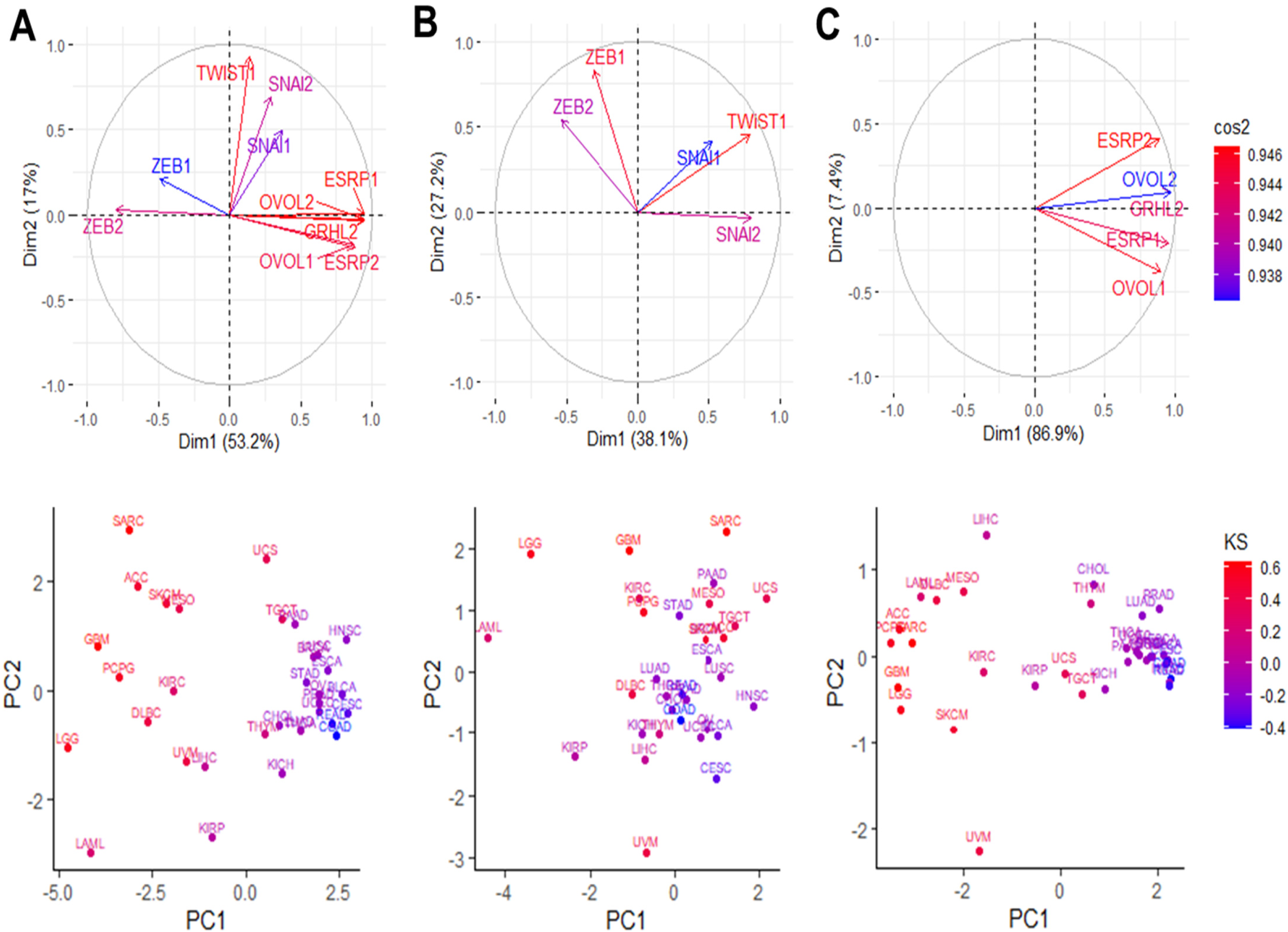
MET factors are the predominant determinant of epithelial-like and mesenchymal-like clusters. Principal component analysis (PCA) of EMT and MET marker expression values. The blue to red color scale represents the contribution of each gene to the segregation of tumor types into epithelial-like and mesenchymal-like clusters. **(A)** PCA of all five EMT and MET markers **(B)** PCA of only EMT markers **(C)** PCA of only MET markers.

**Figure S5:**
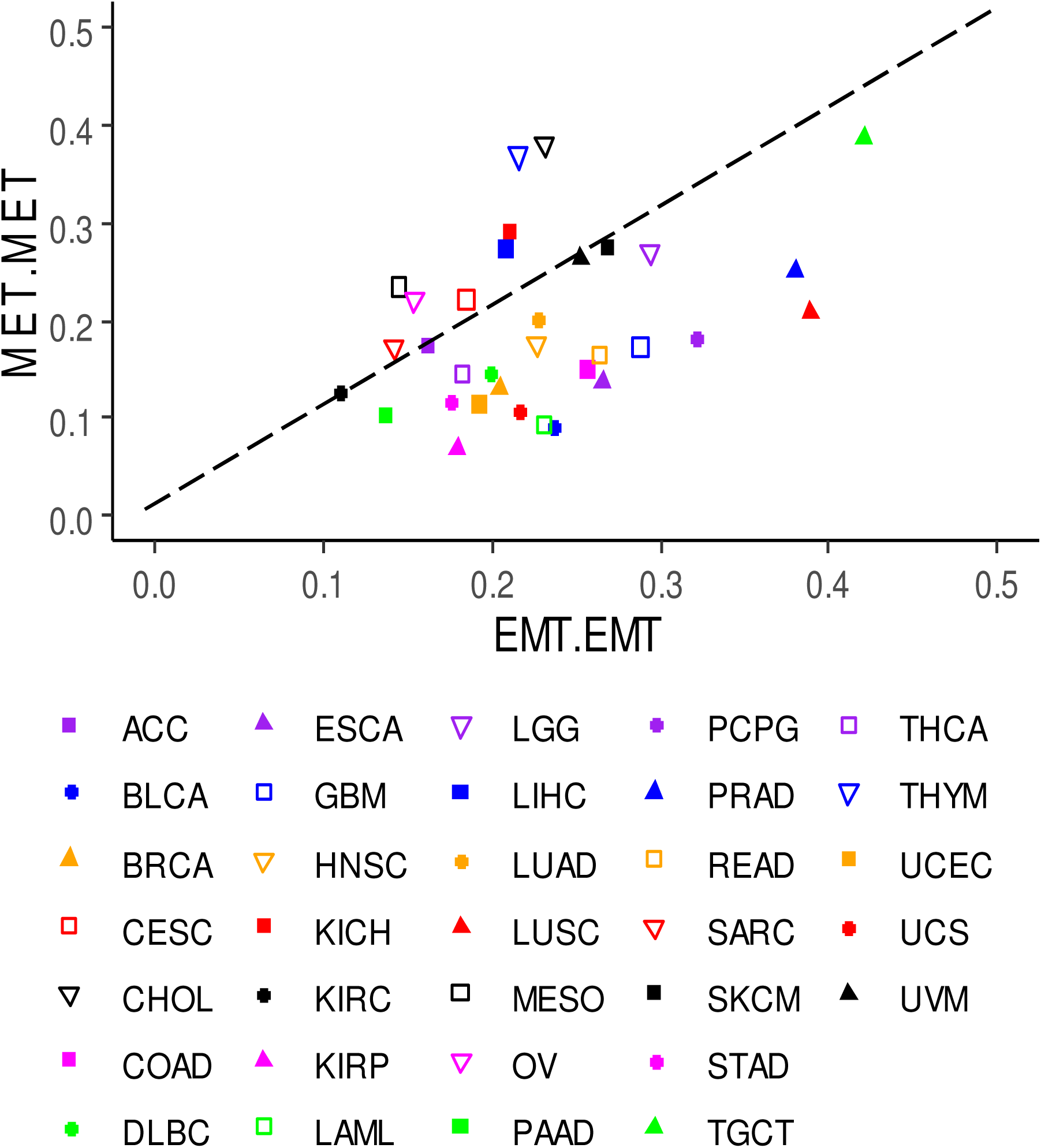
Standard deviation of the correlation of EMT and MET factor expression. Standard deviation of pairwise correlation coefficients between EMT-EMT markers (x-axis) and MET-MET markers (y-axis) across TCGA tumor types

**Figure S6:**
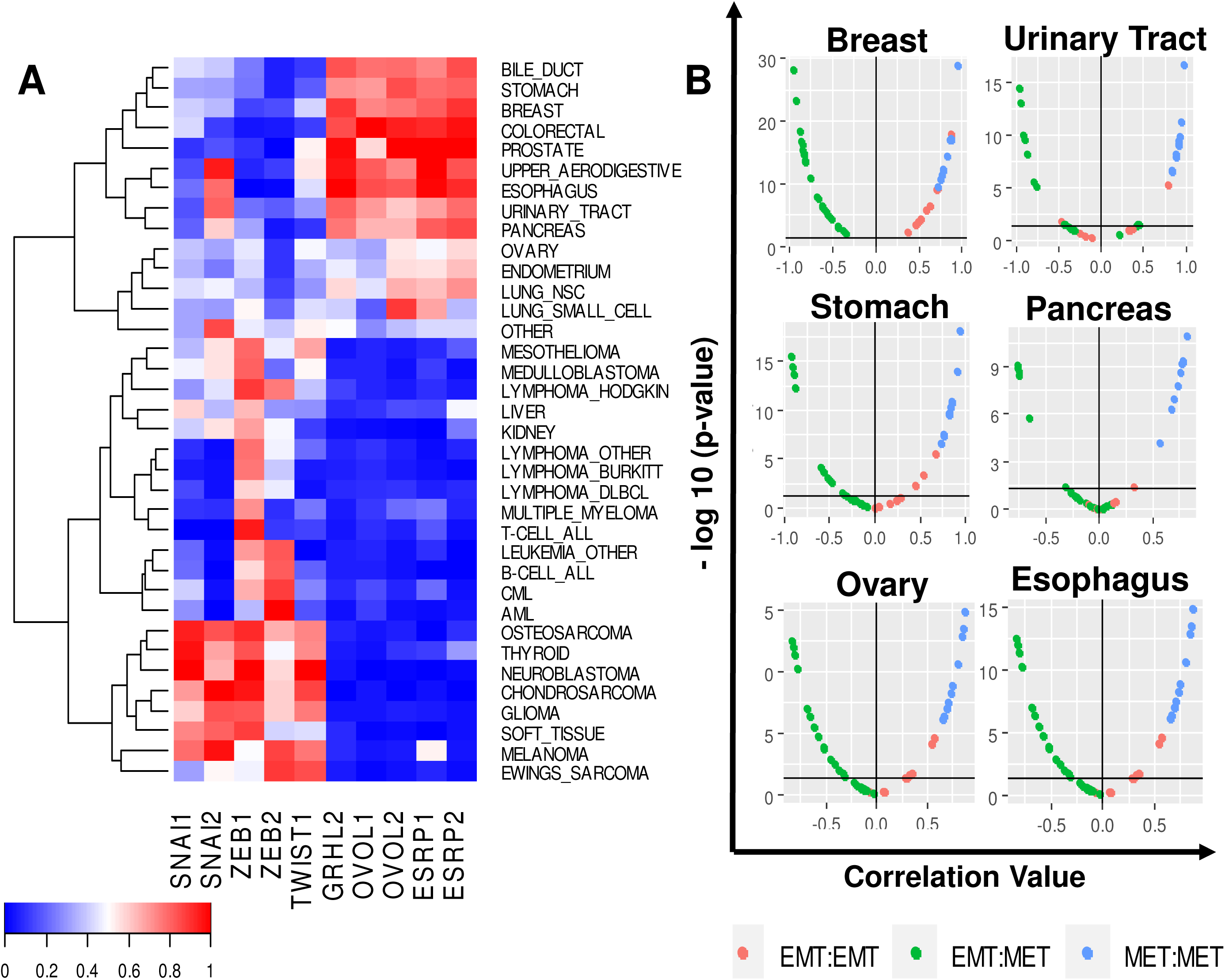
EMT factors are heterogeneously expressed across cancers. EMT and MET marker expression across CCLE tumor types **(A)** Normalized marker expression across tumor types **(B)** Pairwise correlations between expression values of all EMT-EMT, MET-MET, and EMT-MET factor pairs across a subset of tumor types.

**Figure S7:**
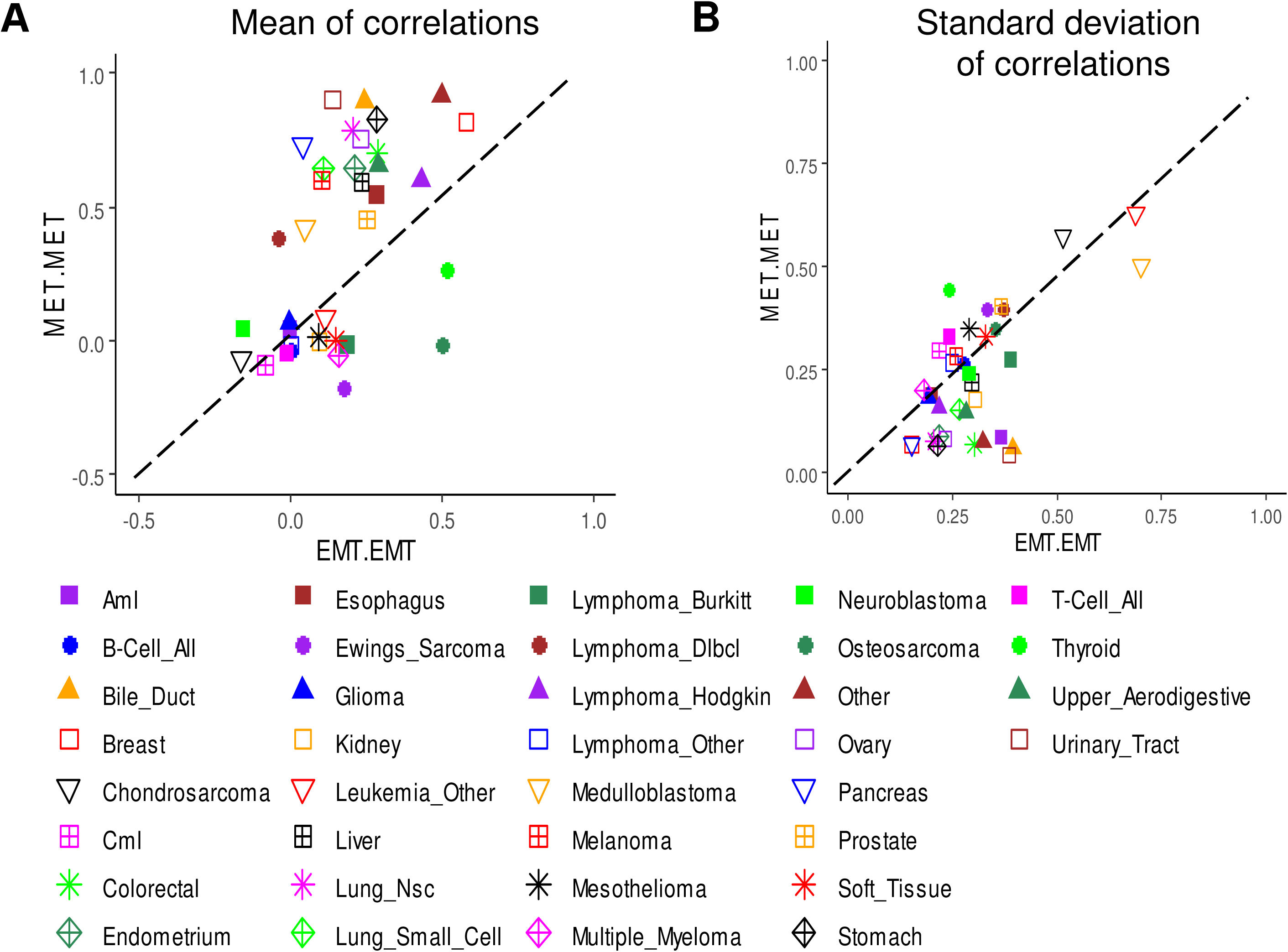
Variability in EMT and MET factor expression. **(A)** Plot of mean pairwise correlation coefficients for all EMT-EMT factor pairs(x-axis) versus mean pairwise correlation coefficients for all MET-MET factor pairs (y-axis) across all CCLE tumor types. **(B)** Standard deviation of pairwise correlation coefficients between EMT-EMT markers (x-axis) and MET-MET markers (y-axis) across CCLE tumor types.

**Figure S8:**
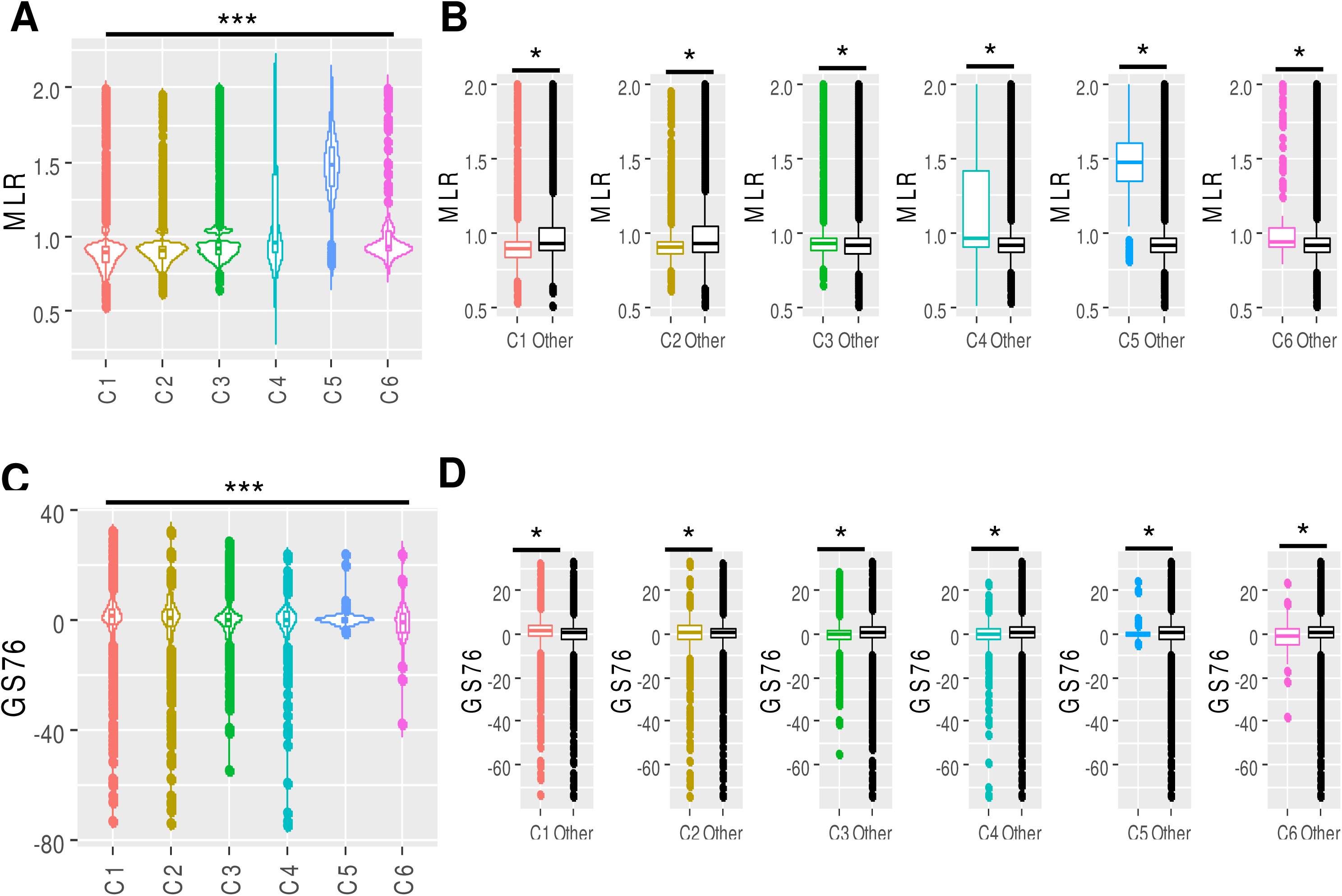
Immune subtypes are associated with EMT scores across scoring algorithms. EMT scores across TCGA immune subtypes; C1 = wound healing, C2 = IFN-γ dominant, C3 = inflammatory, C4 = lymphocyte depleted, C5 = immunologically quiet, C6 = TGF-β dominant **(A)** Plot of calculated MLR scores across all immune subtypes **(B)** Leave-one-out-analysis: pairwise comparison of each cancer immune subtype’s MLR score to the MLR scores of all other immune subtypes **(C)** Plot of calculated 76GS scores across all immune subtypes **(D)** Leave-one-out-analysis: pairwise comparison of each cancer immune subtype’s 76GS score to the 76GS scores of all other immune subtypes

**Figure S9:**
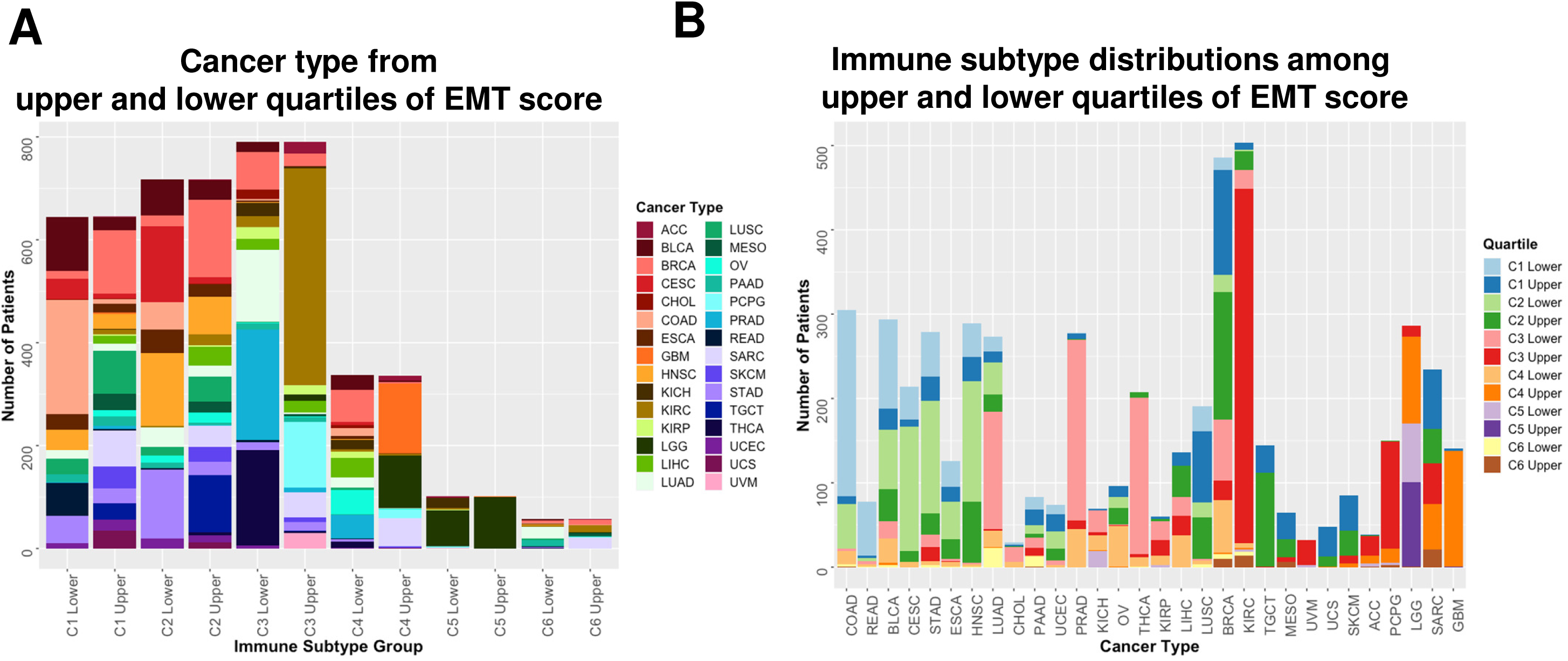
Extremes of EMT score are enriched for specific cancer types across immune subtypes. **(A)** Analysis of TCGA cancer type distribution across the upper and lower EMT score quartiles of all immune subtype groups; COAD = Colon adenocarcinoma, READ = Rectum adenocarcinoma, CESC = Cervical squamous cell carcinoma and endocervical adenocarcinoma, BLCA = Bladder urothelial carcinoma, STAD = stomach adenocarcinoma, ESCA = esophageal carcinoma, UCEC = uterine corpus endometrial carcinoma, LUAD = lung adenocarcinoma, HNSC = head and neck squamous cell carcinoma, PRAD = prostate adenocarcinoma, CHOL = cholangiocarcinoma, PAAD = pancreatic adenocarcinoma, KICH = kidney chromophobe, OV = ovarian serous cystadenocarcinoma, LUSC = lung squamous cell carcinoma, THCA = thyroid carcinoma, BRCA = breast invasive carcinoma, KIRP = kidney renal papillary cell carcinoma, LIHC = liver hepatocellular carcinoma, THYM = thymoma, LAML = acute myeloid leukemia, KIRC = kidney renal clear cell carcinoma, TGCT = testicular germ cell tumors, MESO = mesothelioma, UCS = uterine carcinosarcoma, DLBC = lymphoid neoplasm diffuse large B-cell lymphoma, UVM = uveal melanoma, SKCM = skin cutaneous melanoma, PCPG = pheochromocytoma and paraganglioma, LGG = brain lower grade glioma, SARC = sarcoma, GBM = glioblastoma multiforme **(B)** Analysis of the upper and lower EMT score quartiles of immune subtype across TCGA cancer types

**Figure S10:**
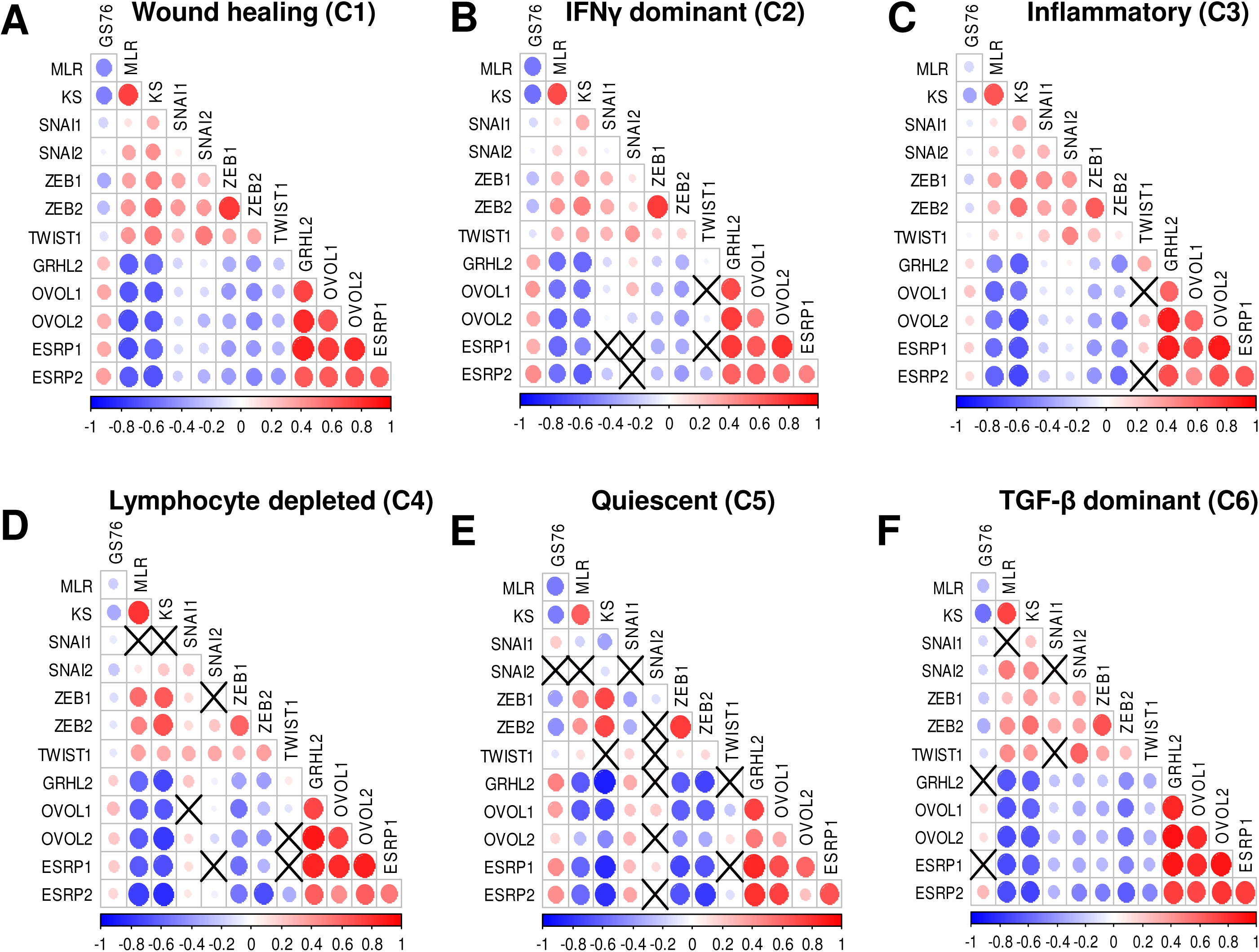
EMT scores correlate with EMT and MET factor gene expression. Correlation between EMT factors, MET factors, and EMT scores across all immune subtypes (A-F)

